# Repurposing of the enhancer-promoter communication underlies the compensation of *Mesp2* by *Mesp1*

**DOI:** 10.1101/2021.06.28.450114

**Authors:** Hajime Okada, Yumiko Saga

## Abstract

Organisms are inherently equipped with buffering systems against genetic perturbations. Upregulation of homologous genes responding to gene loss, termed genetic compensation, is one such buffering mechanism. Recently, a well-conserved compensatory mechanism was proposed: transcriptional adaptation of homologs under the nonsense-mediated mRNA decay pathways. However, this model cannot explain the onset of all compensatory events. We report a novel genetic compensation mechanism operating over the *Mesp* gene locus. *Mesp1* and *Mesp2* are homologs located adjacently in the genome. *Mesp2* loss is partially rescued by *Mesp1* upregulation in the presomitic mesoderm (PSM). Using a cultured PSM induction system, we reproduced the compensatory response *in vitro* and found that the *Mesp2*-enhancer is required to promote *Mesp1*. We revealed that the *Mesp2*-enhancer directly interacts with the *Mesp1* promoter, thereby upregulating *Mesp1* expression upon the loss of *Mesp2*. Of note, this interaction is established by genomic arrangement upon PSM development independently of *Mesp2* disruption. We propose that the repurposing of this established enhancer-promoter communication is the mechanism underlying this compensatory response for the upregulation of the adjacent homolog.

## Introduction

Organisms, especially multicellular organisms, are programmed to develop tissues, organs, and whole bodies. In parallel with these programs, buffering systems against gene network perturbations are inherently present (Tautz, 1992). Genetic compensation is one of these buffering systems. This mechanism is considered to make up for network failure by upregulating homologous genes. Although compensatory events in several types of model organisms have been extensively documented since yeast in 1969 (El-Brolosy & Stainier, 2017), the underlying mechanisms have been described as a consequence of the loss of protein function (O’Leary *et al*, 2013; El-Brolosy & Stainier, 2017) or simply as unknown. Recently, a nonsense-mediated mRNA decay (NMD) pathway-mediated transcriptional adaptation model was proposed. In this model, mutant mRNA bearing a pre-termination codon (PTC) is degraded, and the subsequent pathways modify and activate the chromatin state of the homologous genes of the mutant via fragments of the degraded RNA (Rossi *et al*, 2015; El-Brolosy *et al*, 2019; Ma *et al*, 2019; Serobyan *et al*, 2020). Thus, this mechanism explains the upregulation of homologous genes independent of the downstream protein loss.

Although the NMD-mediated model is widely applicable to genetic mutations that induce genetic compensation, there are exceptional compensatory events. One example is observed in the mutation of mouse *Mesp2,* encoding a transcription factor required for somitogenesis. *Mesp* genes, *Mesp1* and *Mesp2,* are located in a head-to-head orientation on chromosome 7 (Fig 1A), and these genes are co-expressed in the nascent mesoderm and PSM. Replacement of the *Mesp2* locus with *Mesp1* almost completely rescues the *Mesp2* defect (Saga, 1998), suggesting that the functions of MESP proteins are almost identical. Targeted disruption of *Mesp2* induces the compensatory upregulation of *Mesp1*, which partially ameliorates the *Mesp2* defect in the PSM (Takahashi *et al*, 2007). Of note, *Mesp2*-knockout (KO) mice were generated by replacing endogenous genes with exogenous genes, such as *MerCreMer* (Takahashi *et al*, 2007); thus, these mutants do not contain a mutant mRNA-bearing PTC. This strongly suggests that this compensatory response does not rely on the NMD-mediated transcriptional adaptation, but instead on the downstream of MESP2 loss or other compensatory mechanisms.

**Figure 1.**
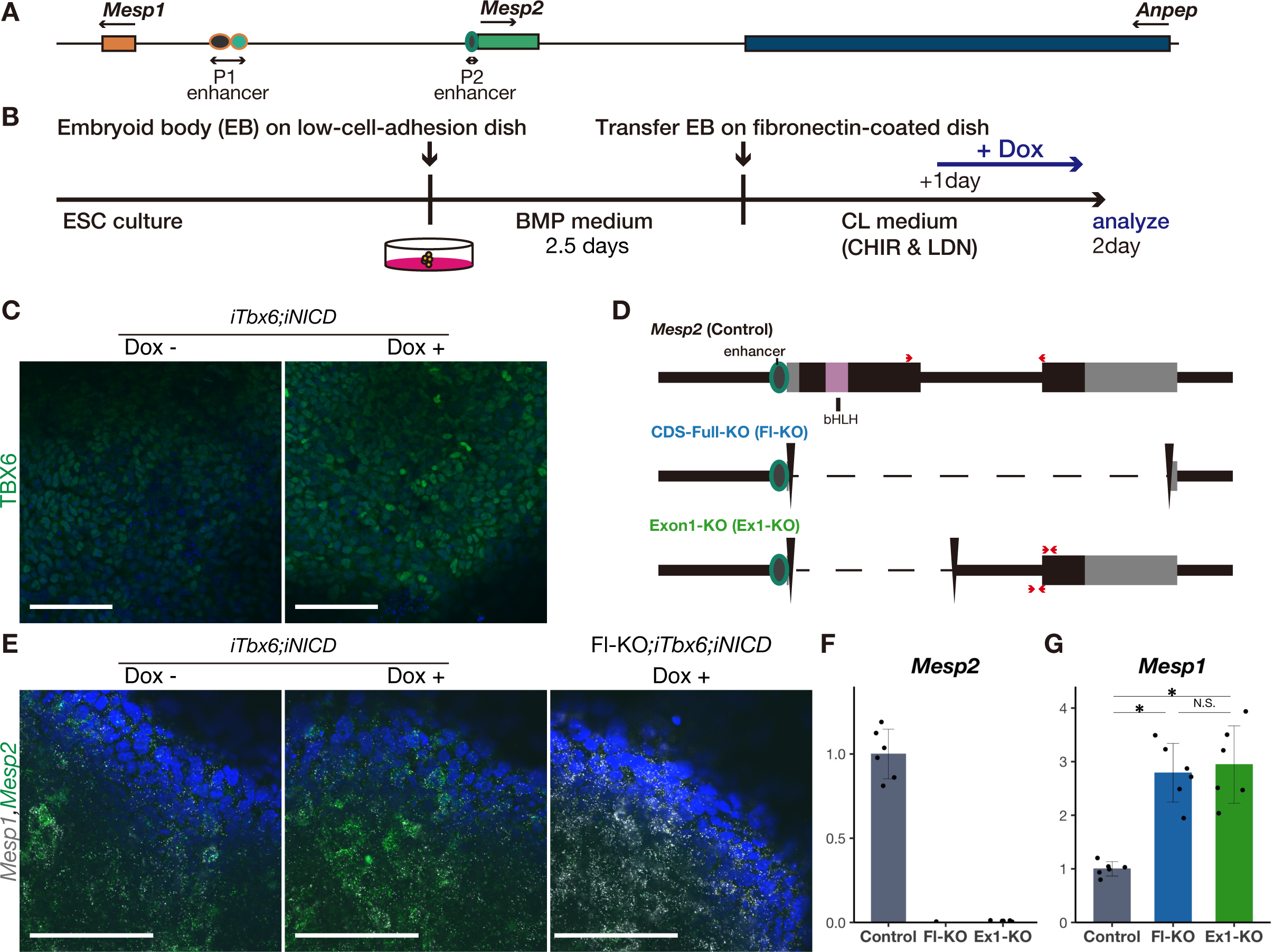
Reproducing the PSM and compensatory response of *Mesp* genes *in vitro.* A Schematic diagram of the genome at the *Mesp2* locus. P1-enhancer and P2-enhancer indicate the *Mesp1*-Enhancer and *Mesp2*-Enhancer, respectively. B Schedule of *in vitro* PSM induction from ES cells for optimal induction of *Mesp2*. C Immunostaining of TBX6 in the *in vitro* PSM of *iTbx6;iNICD* with (right) or without (left) Dox administration. Scale bars: 100 µm. D Schematic diagram of *Mesp2* coding sequence deletions. Red arrows indicate the primers used for qPCR. Primers at *Mesp2* (wild-type), which detect only spliced *Mesp2*, were used for most experiments. Primers at Exon1-KO were used for precursor RNA and either precursor and mature RNA of *Mesp2*, as used in Fig 2A. E *In situ* hybridization for *Mesp1* (white) and *Mesp2* (green) in the *in vitro* PSM of *iTbx6;iNICD* without (left) or with (middle) Dox administration, and *iTbx6;iNICD*;*Mesp2*-CDS-Full-KO upon Dox administration (right). Scale bars: 100 µm. F, G qPCR analysis of *Mesp2* (F) and *Mesp1* (G) in the *in vitro* PSM (n = 6 cultures for each genotype). *P*-values were calculated by the Mann-Whitney U test comparing *iTbx6;iNICD* and *iTbx6;iNICD*;*Mesp2*-KO lines. Data are presented as the mean ± SD. *Asterisk* indicates significant (p < 0.05).

Thus far, enhancer elements have been identified in the intergenic DNA region between *Mesp1* and *Mesp2* (Haraguchi *et al*, 2001; Oginuma *et al*, 2008a). The *Mesp1*-enhancer is located 4k bp distal from *Mesp1* promoter and the *Mesp2*-enhancer is proximal to the *Mesp2* promoter. The activation of either enhancer requires their T-box domains during somitogenesis (Oginuma *et al*, 2008a; Yasuhiko *et al*, 2006). Somites are sequentially formed from the PSM in the anterior to posterior direction. *Mesp1* and *Mesp2* are cyclically upregulated via collaborative Notch signaling and TBX6 in the expected next somite boundary (Morimoto *et al*, 2005, 2006; Oginuma *et al*, 2008b). TBX6 is a T-box transcription factor that binds to either the *Mesp1*-enhancer (Sadahiro *et al*, 2018) or the *Mesp2*-enhancer (Yasuhiko *et al*, 2006, 2008). MESP2 directly induces *Ripply2* (Morimoto *et al*, 2007), and RIPPLY2 then degrades TBX6 through a proteasome pathway leading to the reciprocal transcriptional termination of *Mesp2*, which assures the temporal activation and termination of the *Mesp2* gene (Oginuma *et al*, 2008b; Zhao *et al*, 2015, 2018). When *Mesp2* is disrupted, TBX6 is not sufficiently degraded due to the absence of RIPPLY2 (Sasaki *et al*, 2011; Zhao *et al*, 2015) and may upregulate *Mesp1* via the *Mesp1*-enhancer. Thus, MESP2 protein loss may affect *Mesp1* upstream-regulation, and this breakdown of the negative feedback loop is a possible mechanism of this genetic compensation. In this study, we investigated this possibility and explored a new compensatory mechanism for *Mesp* genes independent of the downstream of MESP2 loss.

## Results

### Establishing an *in vitro* PSM induction system that reproduces the compensatory response of *Mesp1* for *Mesp2* loss

To explore a novel compensatory mechanism for *Mesp* genes during somitogenesis, we took advantage of an established *in vitro* PSM induction system based on the reported protocol (Matsumiya *et al*, 2018). However, we were unable to consistently induce *Mesp2* in this system even though we induced PSM differentiation using the same wild-type (WT) ES cells (Fig. EV1A). The failed induction of *Mesp2* corresponded to lower *Tbx6* and *Notch1* induction, which are essential factors for *Mesp2* induction (Yasuhiko *et al*, 2006) (Fig. EV1A). To reproducibly induce *Mesp2*, we established an ES cell line to induce *Tbx6* and *NICD* upon doxycycline (Dox) administration, named the inducible *Tbx6* and *NICD* (*iTbx6;iNICD*) line. Using this cell line, we induced PSM differentiation with minor optimization of the reported protocol (Fig 1B and see Materials and Methods). Immunocytochemistry (ICC) analysis confirmed the reproducible induction of TBX6 (Fig 1C). We also confirmed the induction of essential developmental genes along with the PSM differentiation process (Fig EV1B and EV1C). Early mesoderm markers, *Brachyury* (*T*) and *Eomes*, were downregulated, and the segmentation clock gene *Hes7*, and segmentation boundary genes *Mesp2* and *Ripply2* were induced by Dox-induced TBX6 and NICD (Fig EV1C). This gene expression pattern supports the efficient induction of PSM in this system.

Next, to examine whether the compensatory response occurred in the cultured system, we deleted the genomic region of *Mesp2* in the *iTbx6;iNICD* ES cell line (the scheme is shown in Fig 1D). Using two coding sequence (CDS)-deleted *Mesp2*-KO lines, CDS-Full-KO (Fl-KO) and Exon1-KO (Ex1-KO) ES cells, we induced PSM differentiation and analyzed the expression level of *Mesp* genes (Fig 1E-G). As previously observed in *Mesp2*-KO mice (Takahashi *et al*, 2007), *Mesp1* was upregulated in the *in vitro* PSM using these two KO lines to an almost equivalent level (Fig 1G). Thus, we succeeded in reproducing the compensatory response in the *in vitro* PSM. Hereafter, we refer to the *iTbx6;iNICD* line and derivative *Mesp2*-KO lines as control and *Mesp2*-KO, respectively.

### The *Mesp2*-enhancer is required for this compensatory response independent of the NMD pathway

The NMD pathway requires mutant mRNA-bearing PTC in the exons before the last exon during the precursor mRNA (pre-mRNA) splicing (Brogna & Wen, 2009). Considering this rule, it is likely that NMD pathways are not activated in *Mesp2*-KO lines because they are lacking almost the entire exon 1 and have no or one exon left (Fig 1D). To clarify this hypothesis that the NMD pathway was not activated in the *Mesp2*-KO lines, we investigated the lack of mRNA degradation after splicing. We examined the expression of pre-mRNA and mature mRNA in the Exon1-KO line, which retains the posterior intronic element and exon 2, using qPCR (primers are shown in Fig 1D). The expression levels of these *Mesp2* transcripts were markedly reduced in this KO line (Fig 2A). This suggests that *Mesp2*-locus transcription itself was suppressed in the Exon1-KO, raising the possibility that this compensatory response occurred independently of the NMD pathway stimulated by PTC-bearing mRNA.

**Figure 2.**
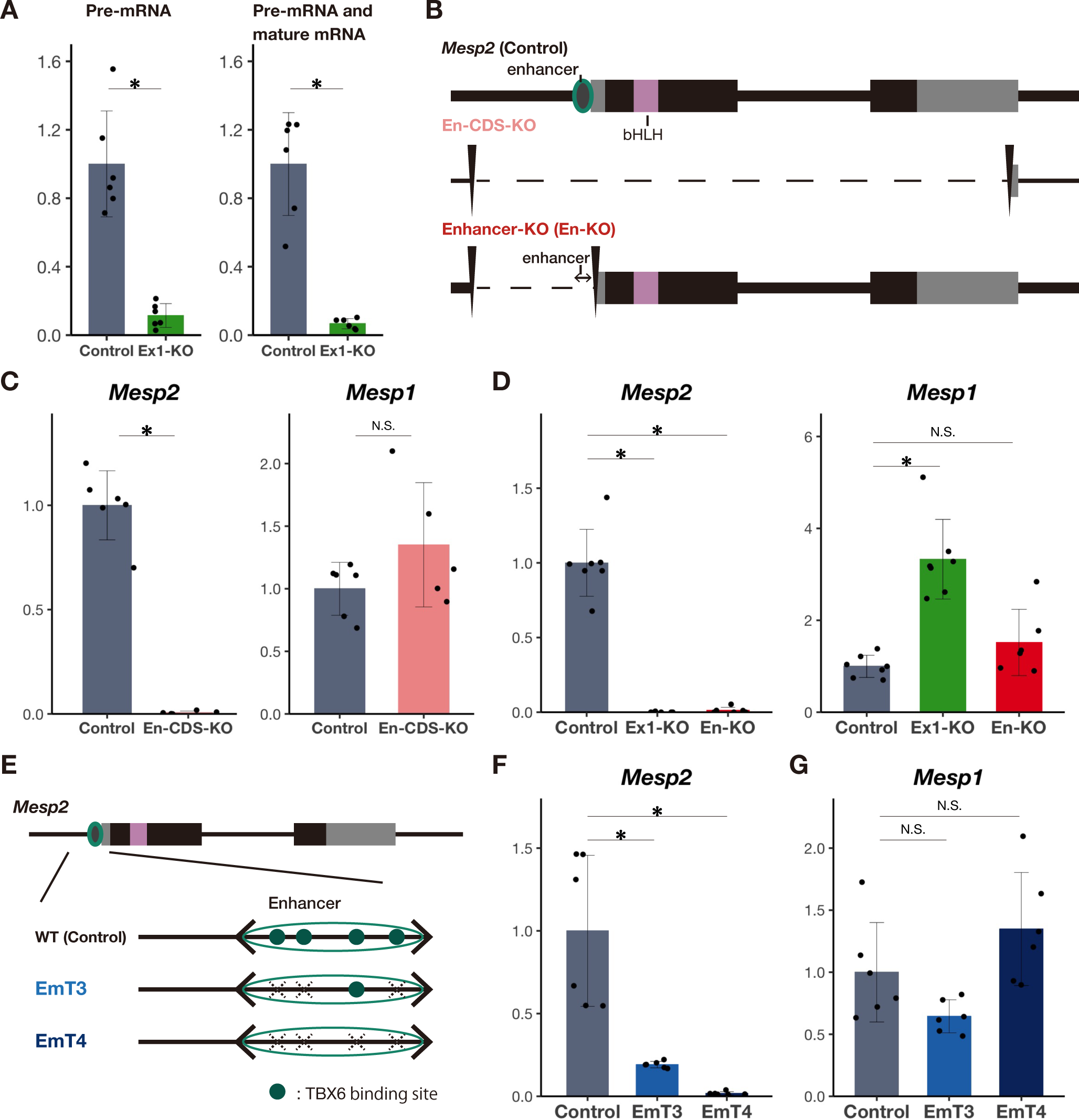
The *Mesp2*-enhancer is required for the compensatory response. A qPCR analysis of precursor RNA (Pre-mRNA) and either precursor or mature RNA of *Mesp2* (n = 6 cultures for each genotype). Used primers are shown in Fig 1D. B Schematic diagram of *Mesp2* genomic deletions of En-CDS-KO and Enhancer-KO. C qPCR analysis of *Mesp2* and *Mesp1* in *in vitro* PSM (n = 5 or 6 cultures for each genotype). *P*-values were calculated by the Mann-Whitney U test comparing *iTbx6;iNICD* and *iTbx6;iNICD*; *Mesp2*-En-CDS-KO. *Asterisk* indicates significant (*p* < 0.05). D qPCR analysis of *Mesp2* and *Mesp1* in *in vitro* PSM (n = 6 to 7 cultures for each genotype). *P*-values were calculated by the Mann-Whitney U test comparing *iTbx6;iNICD* and *iTbx6;iNICD*;*Mesp2*-Exon1-KO or *iTbx6;iNICD*;*Mesp2*-Enhancer-KO. Data are presented as the mean ± SD. *Asterisk* indicates significant (*p* < 0.05). E Schematic diagram of mutations of T-boxes in the *Mesp2*-enhancer. F, G qPCR analysis of *Mesp2* (F) and *Mesp1* (G) in *in vitro* PSM (n = 6 cultures for each genotype). *P*-values were calculated by the Mann-Whitney U test comparing *iTbx6;iNICD* and *iTbx6;iNICD*;*Mesp2*-Enhancer mutant lines. *Asterisk* indicates significant (*p* < 0.05).

The above results led us to hypothesize that the absence of *Mesp2* transcription triggers the compensatory response. To test this hypothesis, we deleted the entire *Mesp2* gene, including both the enhancer and CDS regions, termed En-CDS-KO (Fig 2B). *Mesp2* was depleted in this line, but *Mesp1* was not upregulated (Fig 2C). Thus, the absence of *Mesp2* transcription is not the trigger of the compensatory response and raised another possibility that the *Mesp2*-enhancer is required for this compensation. To test this possibility, we deleted only the *Mesp2*-enhancer region, termed Enhancer-KO (En-KO) (Fig 2B). *Mesp2* transcription was absent in this mutant line, as expected. Consistent with the En-CDS-KO line, *Mesp1* was not upregulated in the Enhancer-KO line (Fig 2D), indicating that the *Mesp2*-enhancer is required for this compensation.

The *Mesp2*-enhancer has four TBX6 binding sites (Fig 2E), and multiple mutations in these TBX6-binding sites reduce the activity of *Mesp2*-enhancer in a stepwise manner (Yasuhiko *et al*, 2008). To address whether activation of the *Mesp2*-enhancer is required for this compensatory response, we disrupted three or four TBX6-binding sites in the *iTbx6;iNICD* line, referred to as EmT3 and EmT4 lines, respectively (Fig 2E). We induced differentiation of EmT3 and EmT4 lines into PSM, and analyzed the expression of *Mesp* genes. As expected, the expression of *Mesp2* was markedly suppressed (Fig 2F) and compensatory upregulation of *Mesp1* did not occur (Fig 2G). Collectively, this indicated that *Mesp2*-enhancer activity is required for this compensation (summarized in Fig EV2).

### The TBX6 increase via MESP2 protein loss may increase *Mesp2*-enhancer activity and the compensatory response

The *Mesp2*-enhancer is required for compensation independent of the NMD pathway. In addition to the NMD pathway, the direct effects of the loss of protein function have been reported as a primary trigger of a compensatory response (El-Brolosy & Stainier, 2017). For example, the disruption of *Rpl22* upregulates its homolog *Rpl22l1*, the expression of which is normally inhibited by *Rpl22* (O’Leary *et al*, 2013). Therefore, breakdown of the negative feedback loop by gene disruption may induce the upregulation of its counterpart homologous gene. In the case of *Mesp*, MESP2 induced by TBX6 induces *Ripply2* (Morimoto *et al*, 2007), and then RIPPLY2 degrades TBX6 through a proteasome pathway (Oginuma *et al*, 2008b; Zhao *et al*, 2015, 2018). Thus, in the *Mesp2*-KO mice, *Ripply2* is not sufficiently induced and TBX6 is not degraded (Sasaki *et al*, 2011; Zhao *et al*, 2015) (Fig 3A). We thus asked whether this breakdown of the negative feedback loop by MESP2 protein loss is involved in this *Mesp2*-enhancer-based compensation. First, to confirm the breakdown of the negative feedback loop, we examined the expression of *Ripply2* in the *Mesp2*-KO ES cell lines. As expected, *Ripply2* expression was reduced (Fig 3B). Consistent with the reduction of *Ripply2*, the protein level of TBX6 was higher in the *Mesp2*-KO lines than in the control, reminiscent of *Mesp2*-KO mice (Fig 3C and EV3A). However, TBX6 was not fully degraded even in control PSM, possibly due to the exogenous induction of TBX6 in this system. On the other hand, the mRNA level of *Tbx6* was comparable between the control and mutants (Fig EV3B). This confirmed the breakdown of the negative feedback loop, especially the TBX6 increase, by MESP2 protein loss in our cultured system.

**Figure 3.**
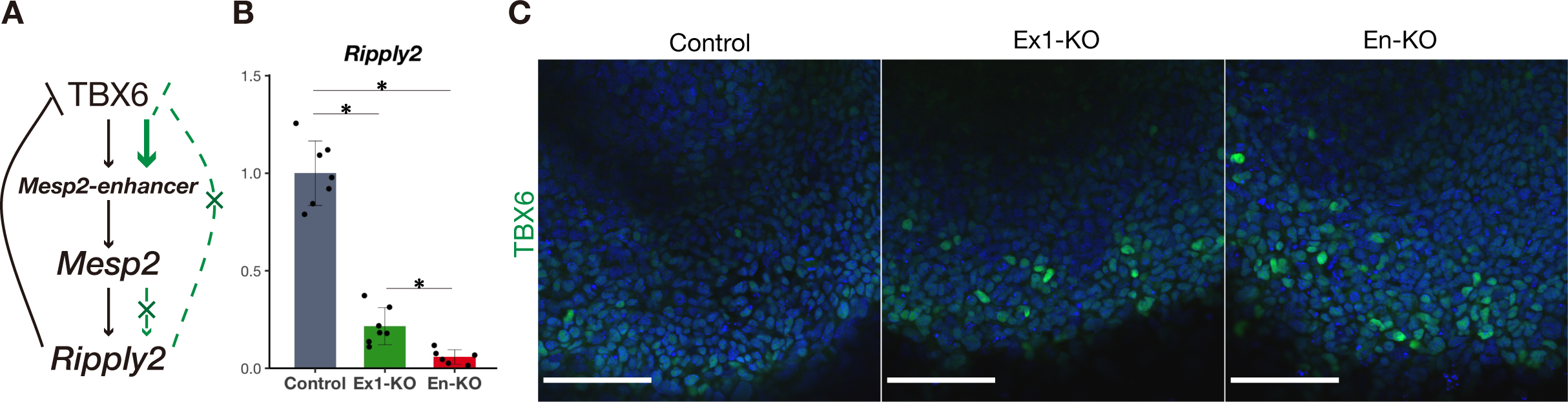
Breakdown of the negative feedback loop by MESP2 loss led to the TBX6 increase. A Schematic diagram of reciprocal regulation of *Mesp2*. Green lines and characters indicate the process of *Mesp2* disruption. B qPCR analysis of *Ripply2* in *in vitro* PSM (n = 6 or 7 cultures for each genotype). *P*-values were calculated by the Mann-Whitney U test comparing *iTbx6;iNICD* and *iTbx6;iNICD*;*Mesp2*-Exon1-KO or *iTbx6;iNICD*;*Mesp2*-Enhancer-KO, or comparing *iTbx6;iNICD*;*Mesp2*-Exon1-KO and *iTbx6;iNICD*;*Mesp2*-Enhancer-KO. Data are presented as the mean ± SD. *Asterisk* indicates significant (*p* < 0.05). C Immunostaining of TBX6 in *in vitro* PSM of *iTbx6;iNICD* (left) and *iTbx6;iNICD*;*Mesp*2-CDS-Exon1-KO (middle), and *iTbx6;iNICD*;*Mesp2*-Enhancer-KO (right) upon Dox administration. Scale bars: 100 µm.

Next, we asked whether this TBX6 increase by MESP2 loss played a role in this enhancer-based compensation. As TBX6 binds T-box binding sites in the *Mesp1*- and *Mesp2*-enhancers (Sadahiro *et al*, 2018; Yasuhiko *et al*, 2008), the higher TBX6 expression in the mutants may function in the compensatory response by promoting *Mesp1*- and *Mesp2*-enhancer activities. However, the deletion or inactivation of the *Mesp2*-enhancer led to no compensatory response of *Mesp1* (Fig 2C, D and G). This indicates that the *Mesp1*-enhancer activity is not affected by the higher level of TBX6 in the absence of the *Mesp2*-enhancer. However, the higher level of TBX6 may increase *Mesp2*-enhancer activity and this compensatory response.

### The *Mesp2*-enhancer promotes *Mesp1* expression in cooperation with the ***Mesp1*-enhancer**

Our study revealed that the *Mesp2*-enhancer plays a central role in compensation and its activity may be promoted by the higher level of TBX6 due to MESP2 loss. The next question is how the *Mesp2*-enhancer regulates this compensation. Enhancers are DNA regulatory elements of gene transcription (Banerji *et al*, 1981). However, the *Mesp2*-enhancer does not regulate *Mesp1,* at least under physiological conditions, because the deletion or inactivation of this enhancer did not lead to the reduction of *Mesp1* (Fig 2C, D, and G). If the *Mesp2*-enhancer promotes the expression of *Mesp1* upon the deletion of the *Mesp2* coding sequence, this enhancer should alter its target from *Mesp2* to *Mesp1*. To test whether the *Mesp2*-enhancer selectively upregulates *Mesp1* or simply affects proximal gene expression, we examined the other gene proximal to *Mesp2*, *Anpep*, which is located adjacent to *Mesp2* on the opposite side of *Mesp1* (Fig 1A). The protein encoded by *Anpep* is an alanyl aminopeptidase that localizes in the plasma membrane and digests peptides, differing in function from MESP proteins. Of note, this non-homologous gene was also upregulated in the condition where *Mesp1* was upregulated (Fig 4A). This supports the idea that the *Mesp2*-enhancer simply affects the proximal genes rather than selectively affecting *Mesp1*.

**Figure 4.**
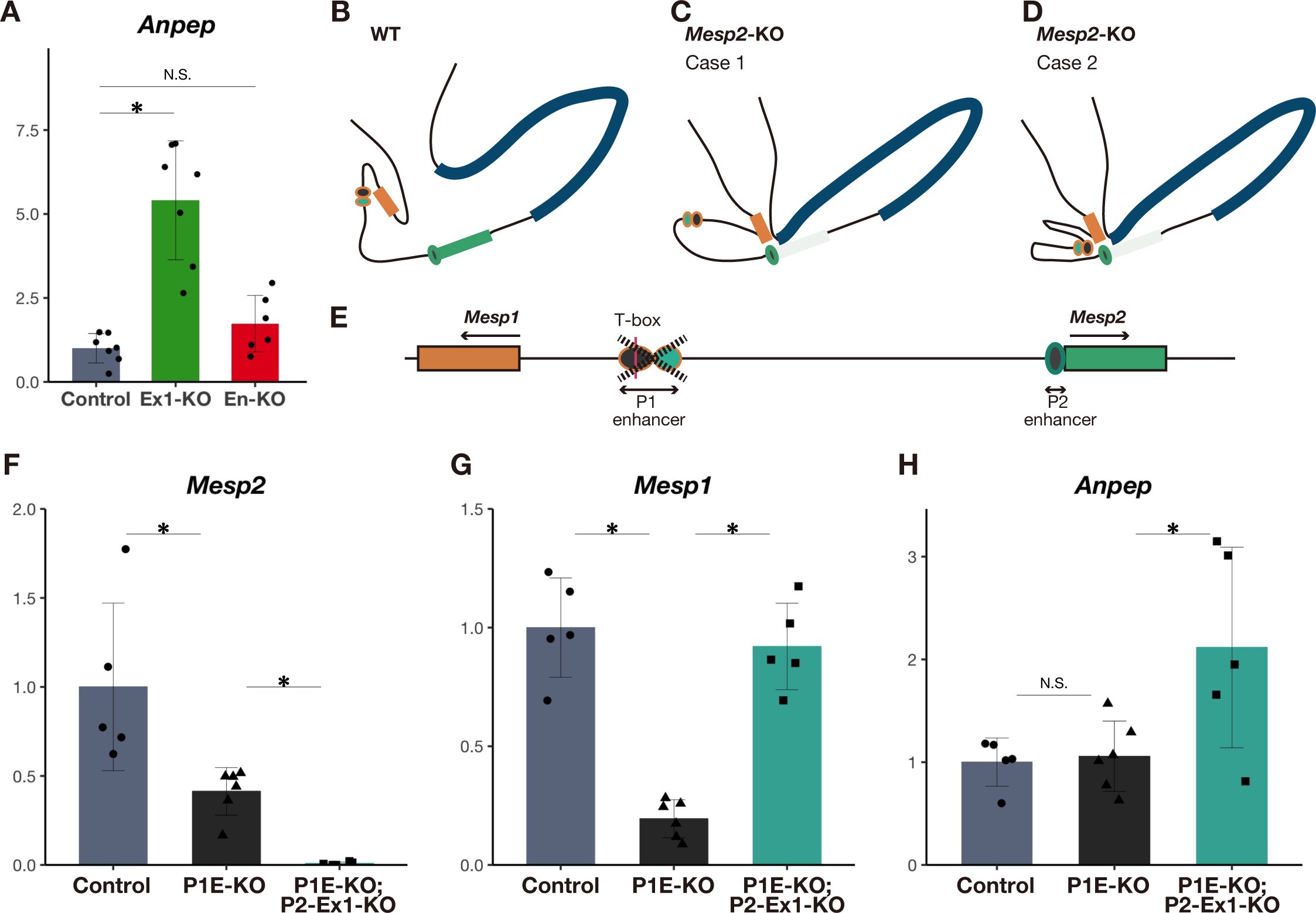
The *Mesp1*-enhancer and *Mesp2*-enhancer cooperatively promote *Mesp1* in the absence of *Mesp2*. A qPCR analysis of *Anpep* in *in vitro* PSM (n = 6 to 7 cultures for each genotype). *P*-values were calculated by the Mann-Whitney U test comparing *iTbx6;iNICD* and *iTbx6;iNICD*;*Mesp2*-Exon1-KO or *iTbx6;iNICD*;*Mesp2*-Enhancer-KO. *Asterisk* indicates significant (*p* < 0.05). B–D The genomic conformation models around the *Mesp2* locus in the presence (B) and absence (C and D) of the *Mesp2* gene. Orange, green, and blue rectangles indicate *Mesp1*, *Mesp2*, and *Anpep*, respectively. Circles indicate enhancers of *Mesp1* (orange line) and *Mesp2* (green line). The *Mesp1*-enhancer interacts with the *Mesp1* promoter in the presence of *Mesp2* (B). If this enhancer interacts with the *Mesp1* promoter in the absence of *Mesp2* (shown as light green), the genome structure should be like case 2 (D). If not, it should be like case 1 (C). E Schematic diagram of genomic deletion of the *Mesp1*-enhancer. Orange-lined black and emerald green circles indicate the *Mesp1*-enhancers activated during early mesodermal and PSM stages and early mesodermal stage only (Oginuma *et al*, 2008a), respectively. The red line indicates the T-box domain. F–H qPCR analysis of *Mesp2* (F), *Mesp1* (G), and *Anpep* (H) in *in vitro* PSM; *iTbx6;iNICD*, *iTbx6;iNICD*;*Mesp1*-enhancer-KO, and additive *Mesp2*-Exon1-KO or *iTbx6;iNICD*;*Mesp1*-enhancer-KO (n = 5 or 6 cultures for each genotype). *P*-values were calculated by the Mann-Whitney U test comparing *iTbx6;iNICD*;*Mesp1*-enhancer-KO and others. Data are presented as the mean ± SD. *Asterisk* indicates significant (*p* < 0.05).

In the presence of *Mesp2*, *Mesp1* must be regulated by the *Mesp1*-enhancer. We hypothesized that the *Mesp1* promoter interacts with the *Mesp1*-enhancer, but not with the *Mesp2*-enhancer under physiological conditions (Fig 4B). On the other hand, in the absence of *Mesp2*, *Mesp1* and *Anpep* promoters should interact with the *Mesp2-*enhancer. In this situation, there are two possibilities regarding the manner by which the *Mesp2*-enhancer interacts with the *Mesp1* promoter. One is that the *Mesp2*-enhancer solely interacts with the *Mesp1* promoter, replaced by the *Mesp1*-enhancer (case 1 in Fig 4C). The other is that the *Mesp2*-enhancer interacts with the *Mesp1* promoter in cooperation with the *Mesp1*-enhancer (case 2 in Fig 4D).

To test these possibilities and elucidate the necessity of the *Mesp1*-enhancer for this compensation, we deleted the *Mesp1*-enhancer region in the *iTbx6;iNICD* cell line (P1E-KO line) (Fig 4E). This deletion led to a marked reduction of *Mesp1* expression and also reduced *Mesp2,* but not *Anpep,* expression (Fig 4F–H), suggesting that the *Mesp1*-enhancer also promotes *Mesp2* expression. To examine the involvement of the *Mesp1-*enhancer in the compensatory response, we further deleted the *Mesp2* coding sequence (shown in Fig 1D) in the P1E-KO line. *Mesp1* upregulation was observed in these lines compared with the P1E-KO line (Fig 4G and EV4), although it did not reach the level observed in the *Mesp2*-KO line containing the intact *Mesp1*-enhancer. *Anpep* was also upregulated in *Mesp2*-KO;P1E-KO more than in P1E-KO, but the basal expression of *Anpep* was not affected by the loss of the *Mesp1*-enhancer (Fig 4H). This indicates that the *Mesp2*-enhancer alone can alter *Mesp1* and *Anpep* expression in the absence of *Mesp2.* However, greater alteration of *Mesp1* may require the *Mesp1*-enhancer (Fig 4G and EV4). The *Mesp1*-enhancer and *Mesp2*-enhancer may cooperatively induce *Mesp1* expression in the absence of *Mesp2*, supporting case 2 (Fig 4D).

### The *Mesp2*-enhancer communicates with the promoters of proximal genes at the PSM stage to alter its targets in the absence of the *Mesp2* coding sequence

The *Mesp2*-enhancer upregulates *Mesp1* and *Anpep* only as a compensatory response (Fig 2D and 4A). We hypothesized that the *Mesp2*-enhancer interacts with the promoters of proximal genes, *Mesp1* and *Anpep,* when it regulates them. To test this hypothesis, we examined the physical interaction of the *Mesp2*-enhancer with the promoters of these genes. We employed engineered DNA-binding molecule-mediated chromatin immunoprecipitation (enChIP) (Fujita *et al*, 2017). We introduced 3×FLAG-dead Cas9 (dCas9) and gRNA, which recognizes the genomic region close to the *Mesp2*-enhancer (Fig EV5A), into the *iTbx6;iNICD* line and the aforementioned *Mesp2*-CDS-Full-KO line. We hereafter refer to these descendent lines as enChIP lines (Fig EV5B). Using enChIP lines, we confirmed the compensatory response in the *in vitro* PSM (Fig EV5C). Then, we pulled down the *Mesp2*-enhancer associated region by FLAG antibody to examine whether the proximal gene promoter regions were associated with the *Mesp2*-enhancer region in the *Mesp2*-KO condition (See Methods).

We hypothesized that the *Mesp2*-enhancer can interact with the proximal gene promoters only in *Mesp2*-KO PSM (Fig 5A and B). To test our hypothesis, we compared the interactions of the *Mesp2*-enhancer before and after PSM induction in the control and *Mesp2*-KO conditions (Fig 5C). As expected, we detected the physical interaction of the *Mesp2*-enhancer with the *Mesp1* and *Anpep* promoters and *Mesp1*-enhancer in the PSM, but not in ES cells (Fig 5D). However, these interactions in the KO PSM were comparable with those in the control PSM. This result, contrary to our hypothesis, indicates that the interaction of the *Mesp2*-enhancer with the proximal gene promoters was established in the PSM, regardless of the presence of *Mesp2*, which negates the assumption of WT genomic conformation in the PSM (Fig 5A and 6A). The interaction frequency of these gene regulatory regions was not increased in the *Mesp2*-KO line (Fig 5D), indicating that the genomic conformation at the *Mesp2* locus was not significantly different between the control and *Mesp2*-KO cells. Thus, this enhancer-promoter communication was differently utilized between the control and *Mesp2*-KO cells. We speculated that the underlying mechanism of this compensation is repurposing of this enhancer-promoter communication to activate different targets: the targets of the *Mesp2*-enhancer are *Mesp2* in the control condition, and *Mesp1* and *Anpep* in the *Mesp2*-KO condition (Fig 6B). The targets of the *Mesp2*-enhancer may be determined depending on the presence of the *Mesp2* coding sequence (Fig 6B).

**Figure 5.**
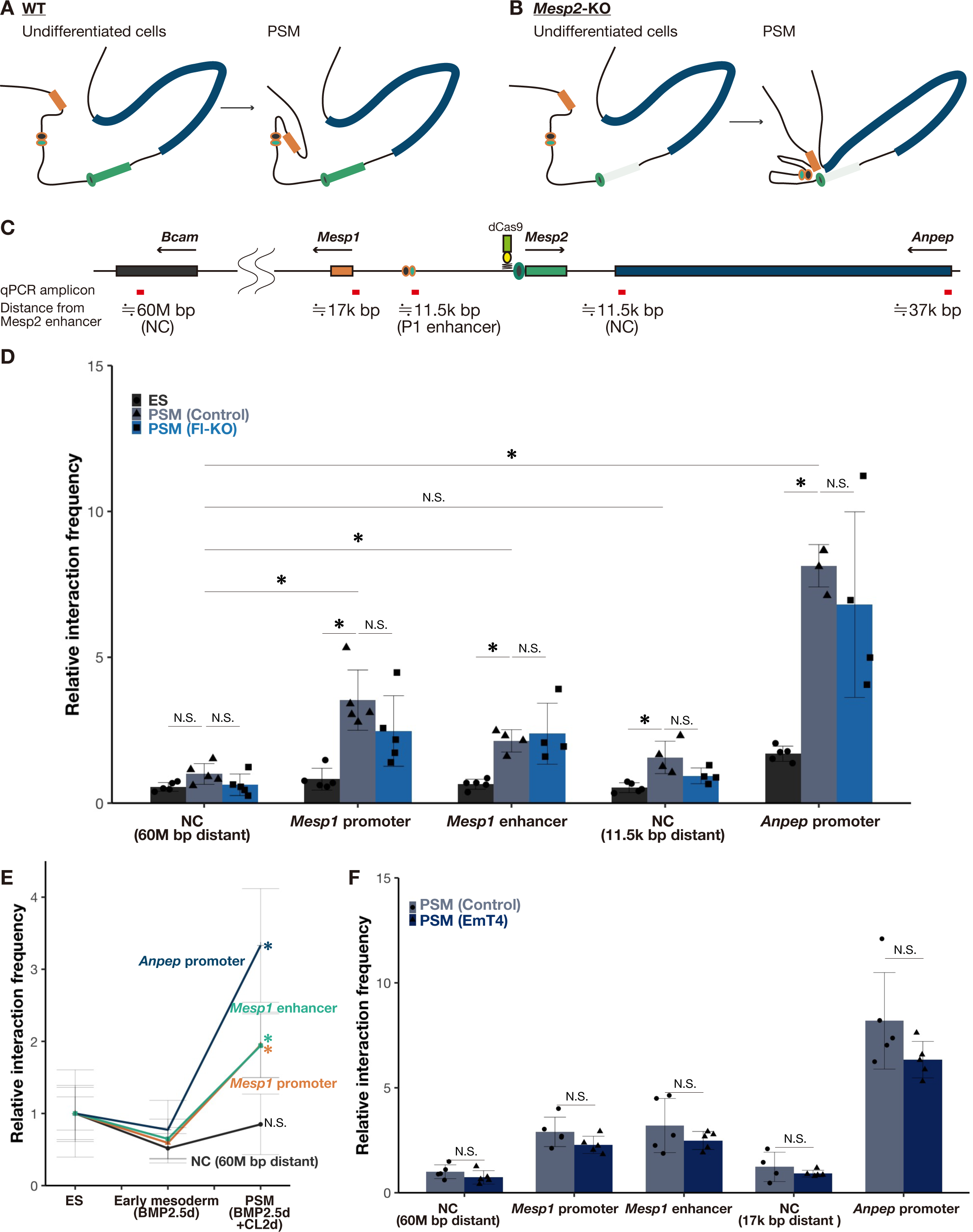
The communication of the *Mesp2*-enhancer with proximal promoters was established in *in vitro* PSM independent of *Mesp2* gene disruption. A, B The genomic conformation models of around the *Mesp2* locus during the differentiation into PSM in the presence (A) and absence (B) of the *Mesp2*gene. Orange, green, and blue rectangles indicate *Mesp1*, *Mesp2*, and *Anpep*, respectively. Circles indicate enhancers of *Mesp1* (orange line) and *Mesp2* (green line). *Mesp2* gene KO is shown as a light green rectangle. C Schematic diagram of the genome around *Mesp2* and *Bcam* regions on chromosome 7. Red bars indicate the examined regions by qPCR for the interaction with the *Mesp2*-enhancer described in Fig 5. D qPCR analysis of physical interactions of genomic regions in IP input and samples from enChIP cell lines (n = 4 or 5 cultures for each sample). Black, grey, and dark blue bars indicate ES cells (control enChIP cell line) and *in vitro* PSM of control and *Mesp2*-CDS-Full-KO enChIP cell lines, respectively. The Y-axis indicates relative interaction frequency, normalized by the value of the 60M-bp region from the *Mesp2*-enhancer in control PSM. *P*-values were calculated by the Mann-Whitney U test comparing regions indicated in this figure. Data are presented as the mean ± SD. *Asterisk* indicates significant (*p* < 0.05). E qPCR analysis of physical interactions of *Mesp1* (Orange) and *Anpep* (Blue) promoters and *Mesp1*-enhancer (Emerald green) and *Bcam* gene-body (Black) regions in IP input and samples from control enChIP cell lines differentiating from ES cells into PSM (n = 5 cultures for each sample). The Y-axis indicates relative interaction frequency, normalized by the value of each promoter interaction in ES cells. *P-*values were calculated by the Mann-Whitney U test comparing ES cells and others in each promoter and the 60M-bp region from the *Mesp2*-enhancer. Data are presented as the mean ± SD. *Asterisk* indicates significant (*p* < 0.05). F qPCR analysis of physical interactions of genomic regions in IP input and samples from enChIP cell lines (n = 5 cultures for each sample). Grey and dark blue bars indicate *in vitro* PSM of control and EmT4 enChIP cell lines, respectively. The Y-axis indicates relative interaction frequency, normalized by the value of the 60M-bp region from the *Mesp2*-enhancer in control PSM cells. *P-*values were calculated by the Mann-Whitney U test comparing regions indicated in this figure. Data are presented as the mean ± SD. *Asterisk* indicates significant (*p* < 0.05).

**Figure 6.**
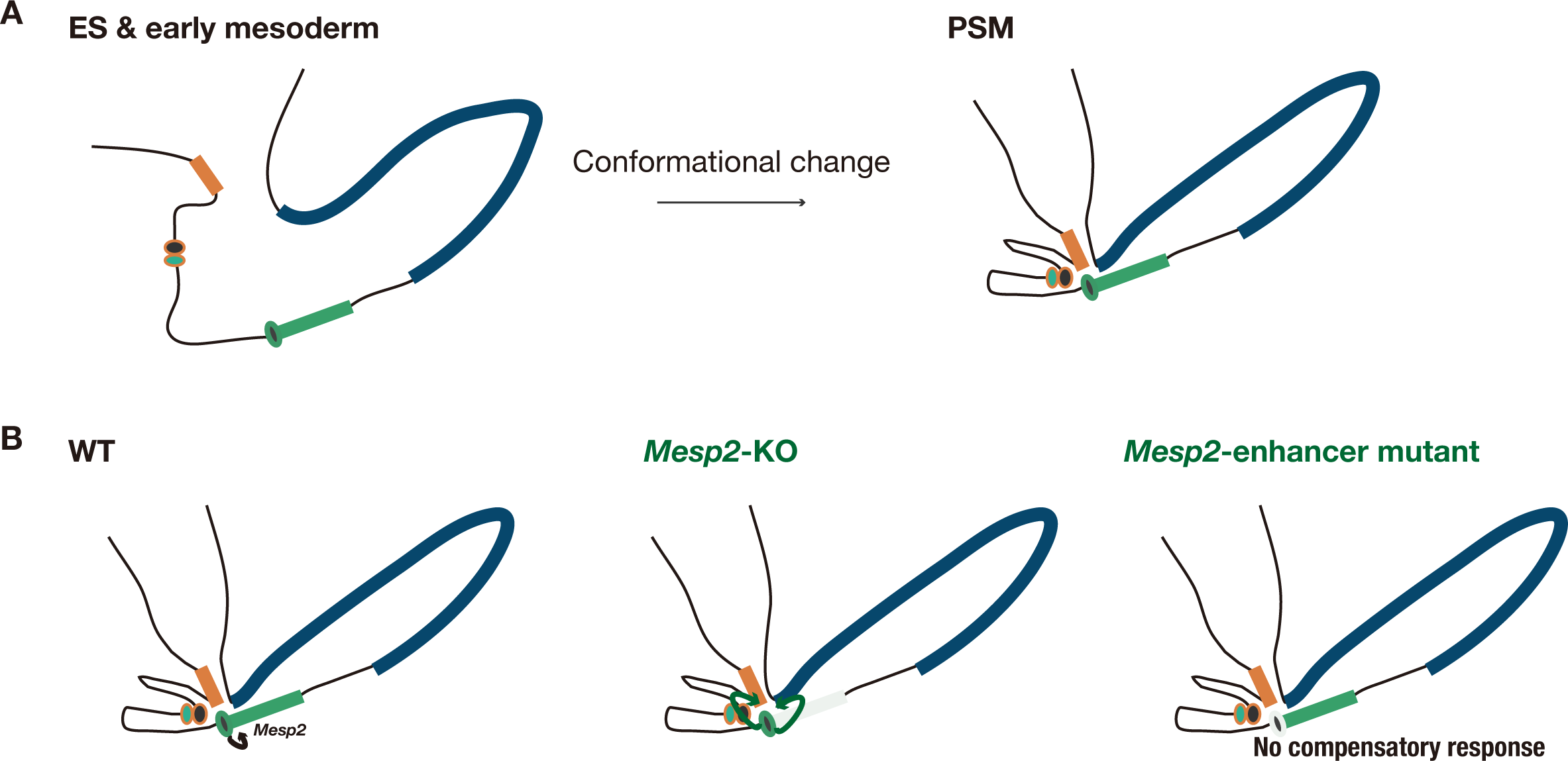
Proposed genome conformation model of genetic compensation among *Mesp2* and proximal genes. The model of genomic conformation around the *Mesp2* locus. Orange, green, and blue rectangles indicate *Mesp1*, *Mesp2*, and *Anpep*, respectively. Circles indicate enhancers of *Mesp1* (orange line) and *Mesp2* (green line). *Mesp2* gene KO and *Mesp2-*enhancer disruption are shown as a light green rectangle and circle, respectively. A In ES cells and early mesoderm cells, the genome around *Mesp2* should be loose (left). In the PSM, the proximal promoters and *Mesp1*-enhancer interact with the *Mesp2*-enhancer (right). B The genomic structure around *Mesp2* is not altered when the *Mesp2* gene is knocked out (shown as a light green rectangle in the middle panel) or the *Mesp2*-enhancer is inactivated (shown as a light green circle in the right panel), compared with the normal condition (left panel). When the *Mesp2* gene is knocked out, this enhancer-promoter communication is repurposed to activate *Mesp1* and *Anpep*. When the *Mesp2*-enhancer is mutated, this enhancer no longer functions and the proximal genes are not activated.

To confirm the specificity of these interactions, we examined 60M bp distant from the *Mesp2*-enhancer region. The *Mesp2*-enhancer communicated with either the *Mesp1* or *Anpep* promoter and the *Mesp1*-enhancer significantly more than this distant region in the PSM. As these interactions are significantly induced in the *in vitro* PSM compared with in ES cells (Fig 5D), the genomic conformation may develop along with PSM differentiation. Regarding its establishment, we examined these interactions in control cells during the differentiation to PSM. As expected, the interactions were established in the PSM, but not in cells treated with BMP medium for two and half days, presumably the early mesodermal cells (Fig 5E). This indicates that the enhancer-promoter communication is established at the PSM differentiation stage (Fig 6A).

The enhancer-promoter communication may play a role in this compensation. Next, we addressed how it is established in the PSM. *Mesp2*, and its upstream regulators, *Tbx6* and *NICD,* were not expressed in ES cells or early mesoderm cells (EV1C). The status of the *Mesp2*-enhancer is active in the PSM but inactive in the ES and early mesoderm cells. The activation status of the *Mesp2*-enhancer may correspond with the existence of this enhancer-promoter communication (Fig 5D and E); therefore, we hypothesized that activation of the *Mesp2*-enhancer induces this enhancer-promoter communication. To address this possibility, we introduced dCas9 and the same gRNA as above into the EmT4 cell line (Fig 2E), which loses the *Mesp2*-enhancer activity due to the lack of TBX6 interaction, and investigated the enhancer-promoter communication in this line. However, these interactions did not disappear (Fig 5F). Thus, neither the activation status of *Mesp2*-enhancer nor TBX6 binding determines this enhancer-promoter communication. The enhancer-promoter communication remained, but the mutated enhancer is no longer able to activate the proximal gene promoters (Fig 2G and 6B). Although TBX6 is important for *Mesp2*-enhancer activity, this factor was not involved in the establishment of this enhancer-promoter communication, and other regulatory factors are expected.

## Discussion

Genetic compensation is a backup mechanism for unexpected disruption of the inherent gene by upregulating its homolog. The underlying mechanism has been ascribed to the induction of downstream signaling in the absence of protein function; however, the molecular mechanisms remain unclear in most cases. Recently, NMD and subsequent pathways were proposed as a mechanism to explain the compensatory response, where PTC-bearing mutant mRNA is involved (Rossi *et al*, 2015; El-Brolosy *et al*, 2019; Ma *et al*, 2019; Serobyan *et al*, 2020). In this study, we investigated the genetic compensation mechanism of *Mesp* genes, which cannot be explained by the NMD-mediated model, and discovered a novel enhancer-based compensatory mechanism.

The most notable finding related to the enhancer-based compensation mechanism is that the communication of the *Mesp2*-enhancer with the *Mesp1* promoter and *Mesp1*-enhancer is established developmentally in the *in vitro* PSM and repurposed for the compensation event (Fig 6). In contrast to the NMD-mediated model, this mechanism requires the establishment of a genomic interaction to evoke the compensation. How such genomic interactions are established and how enhancer targets are determined are points of interest, as discussed below.

### What defines the genomic conformation to induce enhancer-promoter communication?

In this compensatory event, the enhancer-promoter communication (hereafter called E-P communication) was established during PSM development. The critical question is what mechanism regulates the initiation of E-P communication of *Mesp*. The genome is hierarchically organized and shaped into extruding DNA loop structures, topologically associating domains (TADs), mediated by CTCF and cohesin complex proteins at gene boundaries (Dixon *et al*, 2012). Long-range E-P communication, such as crossing TADs, requires domain skipping activity via TAD formation to shorten the inter-domain distance and the distal enhancer to act on the promoter (Yokoshi *et al*, 2020). However, the removal of CTCF or RAD21, a cohesin component, leads to the collapse of TADs, but only modestly affects overall gene expression (Nora *et al*, 2017; Rao *et al*, 2017). E-P communication in more local communities was reported to be facilitated by transcriptional components in some cases (Deng *et al*, 2012; Liu *et al*, 2016). We hypothesized that TBX6 induced this communication because this factor determines the activity of the *Mesp2*-enhancer (Yasuhiko *et al*, 2006, 2008); however, this was not the case (Fig 5F).

TBX6 efficiently binds T-box palindromic sequences as a monomer (White & Chapman, 2005) and also recognizes them as a dimer (Yasuhiko *et al*, 2006). TBX21, also known as T-bet, forms a tight dimer and recognizes T-box DNA elements in the promoter and enhancer of *Ifng* simultaneously, then presumably bridges these elements to induce E-P communication at that locus (Liu *et al*, 2016). Contrary to TBX21, other T-box proteins, TBX1, TBX3, and TBX5, recognize the DNA of two T-boxes as a weaker dimer or two monomers. Thus these T-box proteins were assumed to be unable to form E-P communication based on crystal structure analysis (Liu *et al*, 2016). Collectively, TBX6 likely forms a relatively weaker dimer. It may therefore be reasonable to assume that TBX6 is unlikely involved in this E-P communication based on the DNA recognition manner. Other than proteins of transcriptional components, the transcription at an enhancer, generating noncoding RNA (enhancer-RNA), was reported to establish E-P communication at the immunoglobulin heavy-chain locus (*Igh*) (Fitz *et al*, 2020). These studies suggest that E-P communication can be established in diverse ways and a broader view is required for elucidating the responsible factors.

### The enhancer competition theory is one explanation for the repurposing of enhancer-promoter communication

The next critical question is why E-P communication upregulates *Mesp1* and *Anpep* only when the *Mesp2* gene is disrupted even though the *Mesp2*-enhancer physically interacts with their promoters in the developed PSM. This developmentally regulated E-P communication was also observed in neuron differentiation (Lu *et al*, 2020). In the neuronal cells, E-P communication of essential neural genes is established during the neuronal progenitor phase, and importantly, it does not always correlate with gene activation on a genome-wide scale (Lu *et al*, 2020).

The relative distance between an enhancer and promoters was proposed to determine what promoter is preferentially activated by the enhancer (Fukaya *et al*, 2016). Transcription is not a continuous event, but a bursting event by E-P communication. The proximal promoters compete for enhancer activity, and the closer promoter frequently wins this competition and turns on its transcription (Fukaya *et al*, 2016). As the *Mesp2*-enhancer almost overlaps with the *Mesp2* promoter, the *Mesp2* promoter may always win the competition and dominate the *Mesp2*-enhancer. On the other hand, in the absence of the *Mesp2* coding sequence, this enhancer may function as a distal enhancer for *Mesp1* and *Anpep*. This explains why the *Mesp2*-enhancer is not involved in the regulation of *Mesp1* and *Anpep* under normal conditions, and this enhancer-based compensation should be activated upon *Mesp2*-loss. The switch of an enhancer target without alteration of E-P physical interactions supports this enhancer competition model being applicable to E-P communications that do not accompany gene activation.

The phenomenology of enhancer competition was observed in the target selection by an enhancer. However, the molecular mechanisms, especially what determines the enhancer target, remain unclear. Although E-P communication is important for transcription (Furlong & Levine, 2018), an apparent contradiction was also observed. Live imaging of the *Sox2* locus and *Sox2* distal enhancer element (SCR) revealed that although the *Sox2* locus and SCR can get close, *Sox2* is transcribed regardless of the proximity of SCR (Alexander *et al*, 2019). This suggests that transcription is not simply regulated by physical E-P communication and that an unknown mechanism is involved in the transcriptional regulation by an enhancer. An understanding of general enhancer regulation is necessary to understand the enhancer-based compensation mechanism.

### The involvement of MESP2 protein loss in enhancer-based compensation

We started this study to explore a new compensation mechanism for *Mesp* genes independent of the NMD-mediated pathway and revealed enhancer-based compensation. However, in contrast to the significance of this enhancer-based compensation in our *in vitro* system, *Mesp2*-enhancer mutant mice, corresponding to EmT3 where no compensation occurs in this study (Fig 2E), exhibit the compensatory response (Yasuhiko *et al*, 2008). This strongly suggests another compensation mechanism that functions *in vivo*. What is the difference between *in vivo* and *in vitro*, and what mechanism evokes the compensation *in vivo*? We hypothesize that it is due to the difference in the regulatory mechanism of TBX6 and the activation of the *Mesp1*-enhancer.

TBX6 is degraded through the RIPPLY2-mediated proteasome pathway at the newly forming somite boundary and is never supplied after somite formation in WT mice. Thus, TBX6 is considered to unbind the *Mesp1*- and *Mesp2*-enhancers below a certain threshold, which terminates *Mesp* gene expression (Fig EV6A). On the other hand, in *Mesp2*-KO and *Mesp2*-enhancer mutant mice, TBX6 likely continues to bind the *Mesp1*-enhancer (Fig EV6A). Thus, the different duration of TBX6 binding on the *Mesp1*-enhancer in *Mesp2*-KO and *Mesp2*-enhancer mutant mice compared with that in WT increases the *Mesp1*-enhancer activation period and the amount of *Mesp1* transcripts. On the other hand, in our *in vitro* system, TBX6 and NICD were exogenously induced, and TBX6 was not fully degraded even in control cells (Fig 3C, EV3A, and EV6B). As the *Mesp1*-enhancer has only one TBX6 binding site (Oginuma *et al*, 2008a), this site may always be occupied by TBX6 in the *in vitro* control condition and *Mesp2*-KO condition. There is no more space for TBX6 to bind on the *Mesp1*-enhancer, which may be why the higher level of TBX6 did not alter the *Mesp1*-enhancer activity in *Mesp2*-enhancer KO cells (Fig 2C, D and G).

Of note, the previously reported *Mesp2*-KO mice were generated by replacing *Mesp2* with exogenous genes (Takahashi *et al*, 2007, 2008). If this enhancer-based compensation depends on enhancer competition, this compensation mechanism should not be activated in these *Mesp2*-KO mice. Therefore, instead of the enhancer-based compensation, the prolonged activation of the *Mesp1*-enhancer by TBX6 may be the major compensation mechanism operating *in vivo*. In contrast, we propose that only the enhancer-based compensation mechanism functions in *Mesp1* upregulation in our *in vitro* system. We expect that the mutant mice in which *Mesp2*-enhancer-based compensation occurs will express *Mesp1* more strongly and exhibit a clearer rescue, which is required to evaluate the validity of our interpretation.

### The possible impact of this compensation mechanism

The gene order on the chromosome is not random in eukaryotes (Hurst *et al*, 2004), and adjacent pairs of essential genes for their viability are preferentially conserved through evolution (Pál & Hurst, 2003). Consistent with this, the relative genomic arrangement (synteny) of *Mesp1* and *Mesp2* is evolutionally conserved across zebrafish, mice, and humans. The function and location of the *Mesp2*-enhancer are also well conserved across mice and fish (Terasaki *et al*, 2006; Yasuhiko *et al*, 2008). Moreover, gene pairs that have mutual genomic interaction are highly conserved (Sandhu *et al*, 2012). Collectively, this implies a conserved genomic interaction and compensatory mechanism at this *Mesp* locus across species.

Gene duplications can supply raw materials for evolution (Zhang, 2003; Taylor & Raes, 2004). In addition, gene loss can be a pervasive source of genetic change that drives evolution (Olson, 1999; Albalat & Cañestro, 2016), including some cases of beneficial effects by gene loss (Sharma *et al*, 2018). The prerequisite of the fixation of gene loss in an organism is that the gene is dispensable. The key question is how genes can become dispensable. A study using *Saccharomyces cerevisiae* reported that gene dispensability is predominantly backed up by the transcriptional adaptation of homologs, known as genetic compensation (Kafri *et al*, 2005). Genetic compensation can confer cellular fitness for the survival of an organism; thus this enhancer-based compensation mechanism can be one of the mechanisms making the gene dispensable. Future studies will mine homolog pairs exhibiting this compensatory response other than *Mesp* genes and elucidate generalities. It may be possible to predict which genes are removed and future evolution by combining genome-wide interaction maps such as Hi-C or ChIA-PET data.

## Materials and Methods

### ES cell culture and establishment of modified ES cell lines

ES cells were maintained with feeder cells in ES medium (Yagi *et al*, 1993). Several modified ES cell lines were established by transfection using Lipofectamine 2000. To establish the Tet-inducible *Tbx6*-IRES-*NICD* expression system (*iTbx6;iNICD* cell line), a piggyBac transposon system was used as previously described (Li *et al*, 2013). pBase, CAG-promoter-driven rtTA, and pPB-CMV-*Tbx6*-IRES-*NICD* vectors were transfected into the TT2 ES cells (Yagi *et al*, 1993). The ES cell line was selected using neomycin. To generate several *Mesp2* deletion and *Mesp1*-enhancer deletion lines, gRNAs were transiently introduced into the *iTbx6;iNICD* ES cell line. To introduce point mutations in the *Mesp2*-enhancer, a gRNA and template DNA harboring point mutations in the *Mesp2*-enhancer were transiently introduced into the *iTbx6;iNICD* ES cell line. To detect the physical interaction of the *Mesp2*-enhancer, a piggyBac transposon system was also used. U6-promoter-driven gRNA that recognizes the close DNA region to the *Mesp2*-enhancer (shown in Fig EV5A) and CAG-promoter-driven 3×FLAG-dCas9 vectors were continuously introduced into *iTbx6;iNICD*, *iTbx6;iNICD;Mesp2-*CDS-Full-KO, and *iTbx6;iNICD;*EmT4 ES cell lines, which were selected using puromycin. All gRNAs were designed using CRISPRdirect (Naito *et al*, 2015). gRNA sequences are listed in Table 1.

**Table 1.**
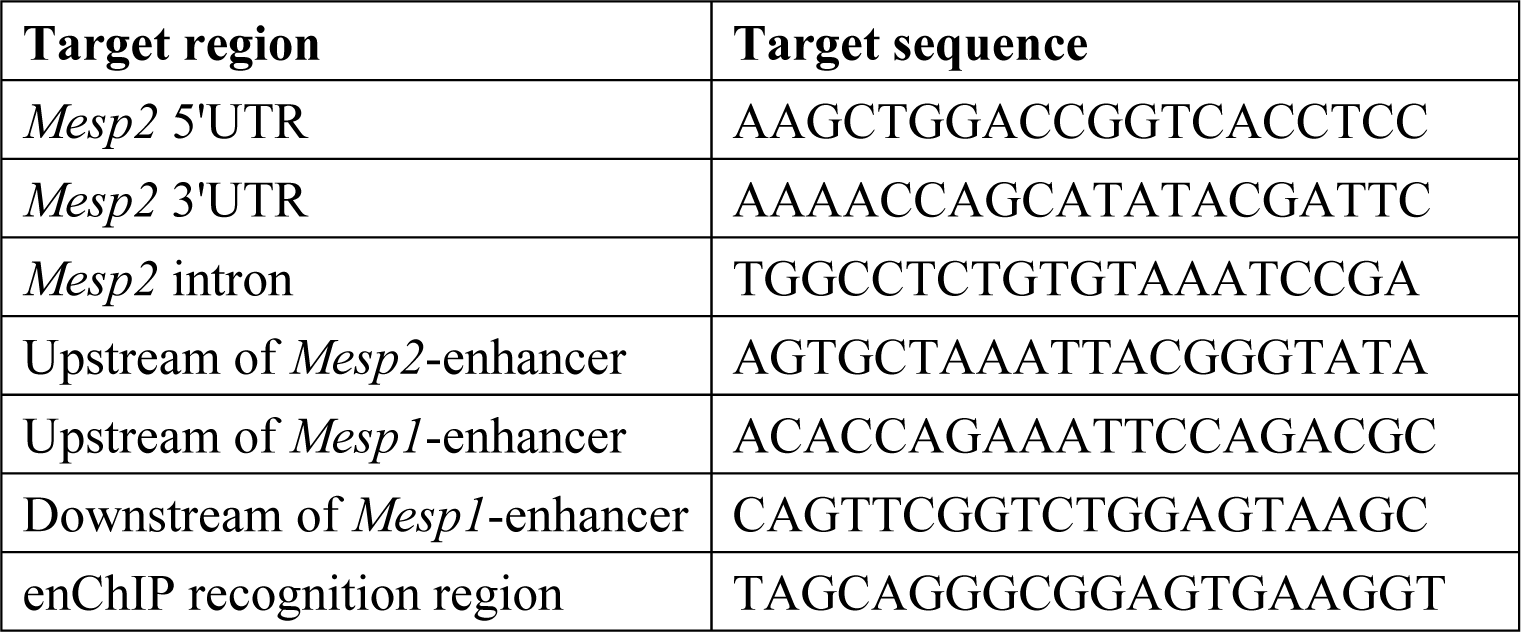
Target sequences of gRNA used in this study.

### *In vitro* PSM induction

For PSM induction, we utilized the method previously described (Matsumiya *et al*, 2018) with some modifications. Briefly, the feeder cells were depleted and ES cells were cultured on gelatin–coated culture dishes for two days before inducing PSM differentiation. One or three thousand (only for enChIP experiments) ES cells per well in low-cell-adhesion 96-well plates with U bottoms were first cultured in DMEM/F12 medium supplemented with N2B27 reagent, 1% Knock-out Serum Replacement (KSR), 0.1% bovine serum albumin, 2 mM L-glutamine, 1 mM nonessential amino acids, 1 mM sodium pyruvate, 10 units/ml of penicillin, 10 mg/ml of streptomycin, and 10 ng/ml of BMP4 (BMP medium) for 2.5 days, which was 2 days in the original paper. Cells were then transferred and cultured on human fibronectin-coated dishes with DMEM medium supplemented with 15% KSR, 2 mM L-glutamine, 1 mM nonessential amino acids, 1 mM sodium pyruvate, 10 units/ml of penicillin, 10 mg/ml of streptomycin, 0.5% DMSO, 1 µM CHIRON99021, and 0.1 µM LDN193189 (CL medium) for 2 days. To induce the Tet-inducible gene expression, 1 mg/mL of doxycycline was added into the medium at CL medium day 1 for one day.

### Visualization of protein and RNA

Immunostaining for cultured cells was performed on cover glasses coated with human fibronectin. Cells were fixed by 4%PFA on ice for 10 min, blocked using 3% FBS, followed by incubating with rabbit-anti-TBX6 (1/200) (Yasuhiko *et al*, 2008) at 4°C overnight and then incubated with Alexa Fluor 488-conjugated anti-rabbit antibody (1/800, Life Technologies, Oregon, USA). For *in situ* hybridization of the mouse *Mesp1* and *Mesp2* mRNA, we used the ViewRNA Cell Plus Assay Kit (Affymetrix, no. TFA-88-19000-99) according to the manufacturer’s instructions. Samples were observed using FluoView FV1200 laser scanning confocal microscopy (Olympus).

### Gene expression analysis by real-time quantitative PCR

Gene expression analysis was performed by real-time quantitative PCR (qPCR). Total RNA was isolated from individual samples using TRIzol reagent (Invitrogen), treated with DNase I (Invitrogen), and then used for cDNA synthesis with Superscript III or IV (Invitrogen) and oligo-dT primer. qPCR analyses were performed using the Thermal Cycler Dice Real Time System (Takara) or C1000 touch Thermal Cycler and CFX Real-Time PCR Detection System (Bio-Rad) with KAPA SYBR FAST Universal 2X qPCR Master Mix (Kapa Biosystems) with optimized concentrations of specific primers. Gapdh was used as the internal control. Primer sequences are listed in Table 2.

**Table 2.**
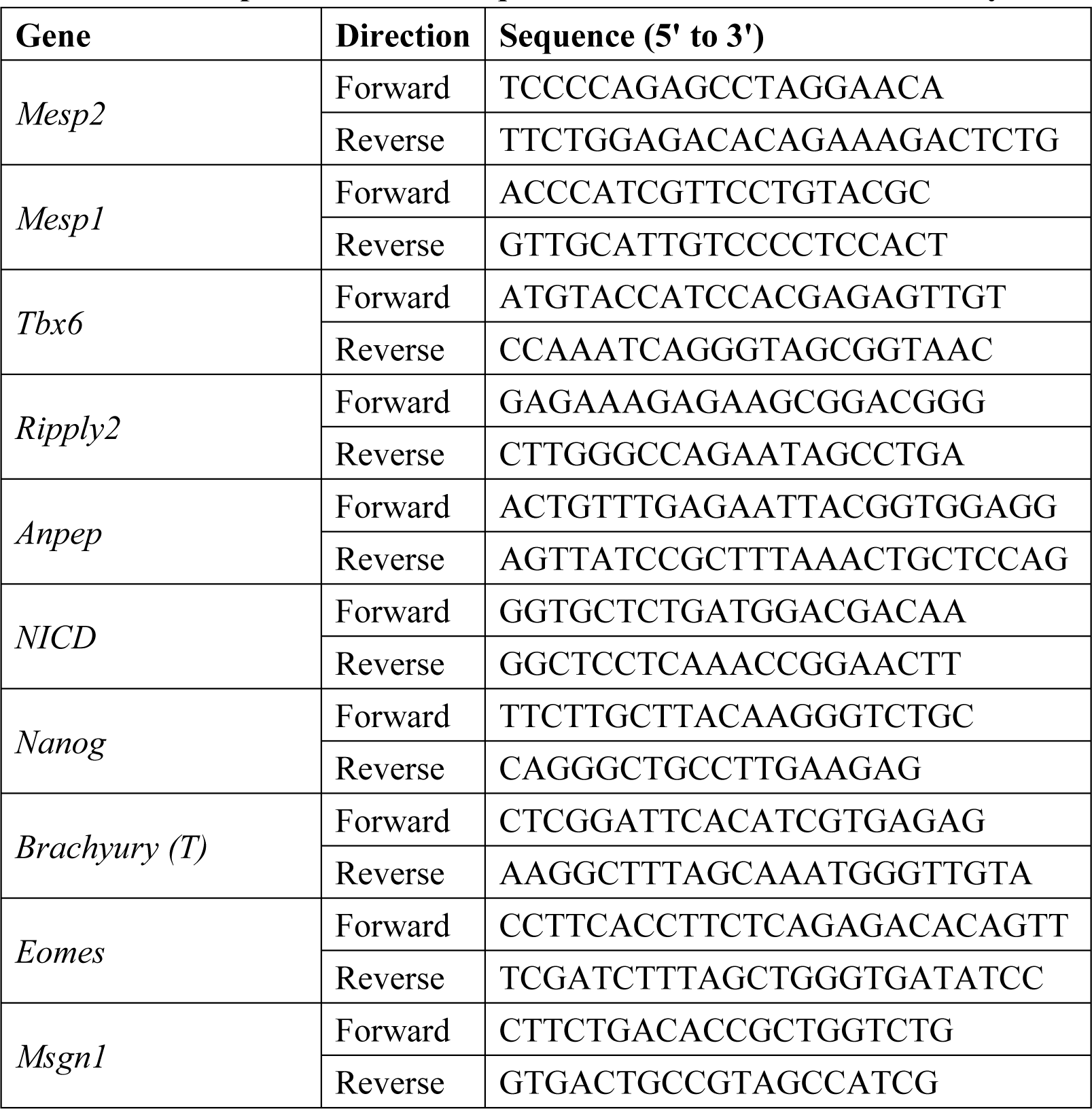
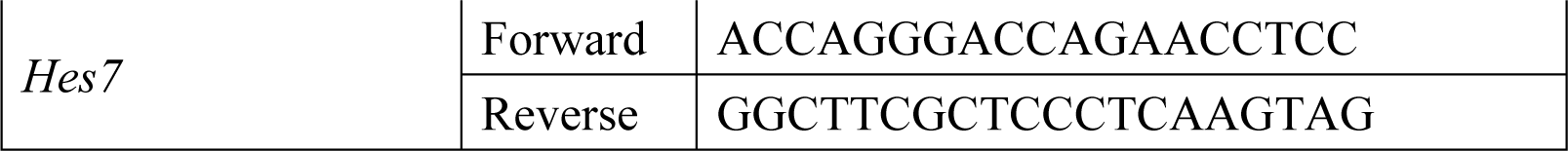
List of primers used for qPCR to detect mRNA in this study.

### Physical interaction assay by engineered DNA-binding molecule-mediated chromatin immunoprecipitation (enChIP)

To detect physical interactions of the *Mesp2*-enhancer with the genome regions of interest, we employed engineered DNA-binding molecule-mediated chromatin immunoprecipitation (enChIP). Cells used for enChIP analyses were generated as mentioned in “ES cell culture and modified ES cell lines establishment.” The procedure of enChIP analysis was described previously (Fujita *et al*, 2017). Briefly, for sample preparation, ES cells 48 hours after seeding 2.0 × 10^5^ cells without feeders on a gelatin-coated 60mm-dish, and 120 wells of low-cell-adhesion 96-well plates during the course of PSM differentiation that started with 3000 cells per well were fixed with 1% formaldehyde at 37 °C for 5 min. Collected samples were lysed and chromatin fractions were extracted. Chromatin fractions were sonicated and immunoprecipitated by Anti-FLAG M2 Magnetic Beads affinity isolated antibody (Sigma, no. M8823). Fragment DNA without immunoprecipitation (input DNA) and immunoprecipitated fragment DNA were used for the subsequent qPCR analyses described in “Gene expression analysis by real-time quantitative PCR.” The occupancy of associated DNA with the *Mesp2*-enhancer was calculated by dividing the immunoprecipitated fragment DNA amount by the input DNA amount. All the data were normalized by immunoprecipitation efficiency estimated by the pull-down efficiency of the *Mesp2*-enhancer fragment. Primer sequences are listed in Table 3.

**Table 3.**
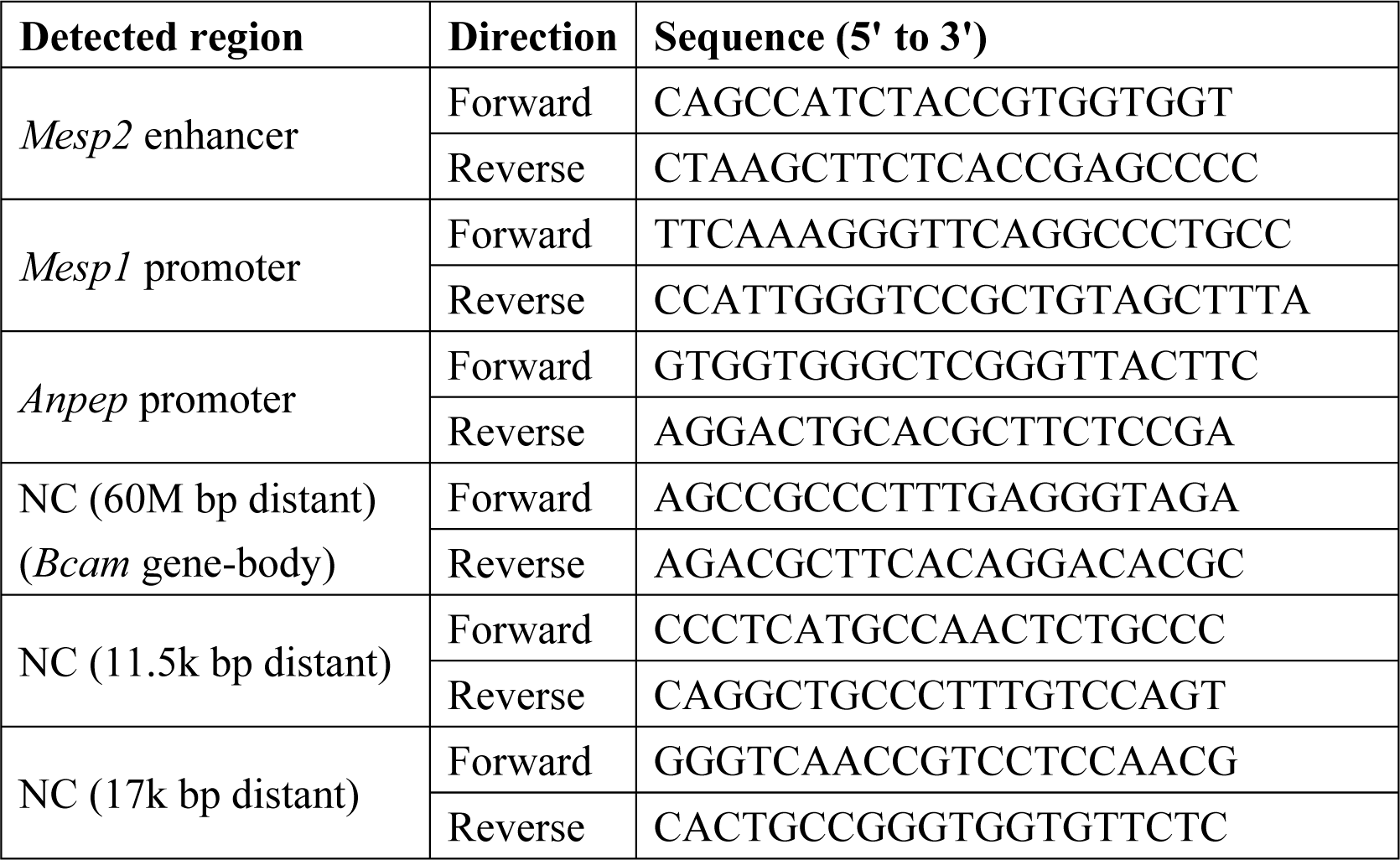
List of primers used for qPCR to detect genomic DNA in this study.

### Statistical analysis

Data are expressed as the mean ± S.D. The Shapiro-Wilk test was used to assess the normality of distribution of investigated parameters and significant differences were tested using the unpaired two-tailed Mann-Whitney U test. Statistical analyses were performed using the R v.3.6.1 software. Differences were considered significant at *p* < 0.05.

## Acknowledgments

We thank Rieko Ajima, Yuzuru Kato, Takamasa Hirano, and Naoko T Fujito (National Institute of Genetics, Japan) for their critical advice and discussions. We thank Makoto Kiso, Masahumi Muraoka, Noriko-Sakurai Yamatani, and Akihiro Maeno (National Institute of Genetics, Japan) for technical support. We thank Danelle Wright (National Institute of Genetics, Japan) for editing this manuscript. We thank Dr. Hitoshi Niwa for providing us piggyBac vectors and Dr. Yukuto Yasuhiko for providing the anti-TBX6 antibody.

This work was supported by the Japan Society for the Promotion of Science KAKENHI grants 19K16152 (to HI) and NIG Postdoctoral Research Fellow grant 2018 (to HI).

## Author contributions

HO, Conception and design, Acquisition of data, Analysis and interpretation of data, Drafting the article, Contributed unpublished essential data or reagents; YS, Conception and design, Analysis and interpretation of data, Drafting the article. All authors reviewed the results and approved the final version of the manuscript.

## Expanded View Figure legends

**Expanded View Figure 1.**
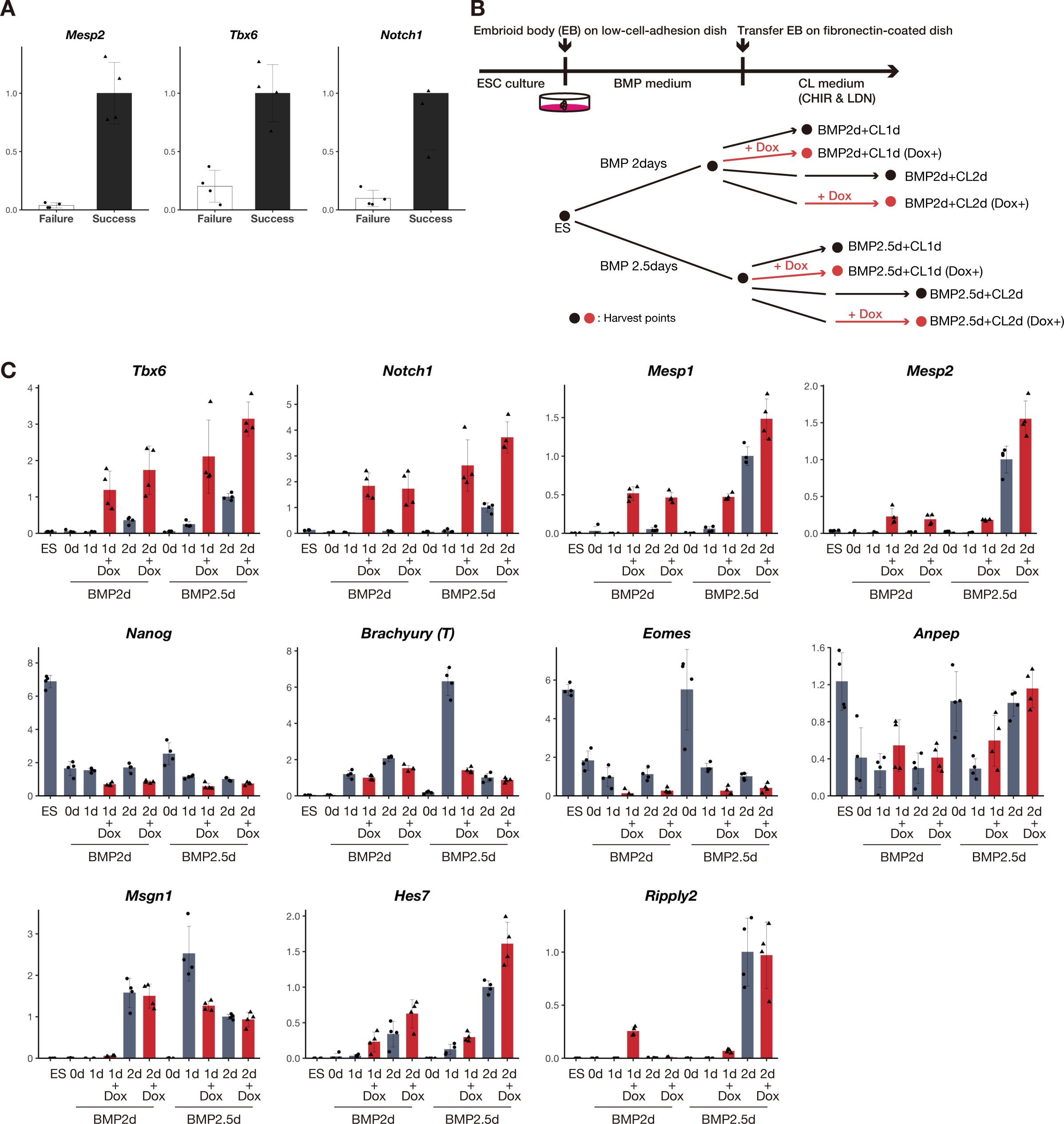
Investigating the expression of developmentally essential genes during PSM development *in vitro.* A qPCR analysis of *Mesp2*, *Tbx6,* and *Notch1* in *in vitro* PSM using wild-type (WT) ES cells. Failure and Success indicate failed and successful PSM induction using the same WT ES cells (n = 4 cultures for each sample). B Schedule of *in vitro* PSM induction from ES cells to cell harvest to analysis. Dox administration is indicated in red. The duration of Dox administration was 24 hours in every set. Red and black circles are harvest time points with or without Dox administration, respectively. C qPCR analysis of representative developmental genes in *iTbx6;iNICD* during PSM induction with (red bars) or without (grey bars) Dox administration (n = 4 cultures for each sample). *Nanog* is a pluripotency marker; *Brachyury* (*T*) and *Eomes* are early mesoderm markers; *Msgn1* and *Hes7* are PSM markers.

**Expanded View Figure 2.**
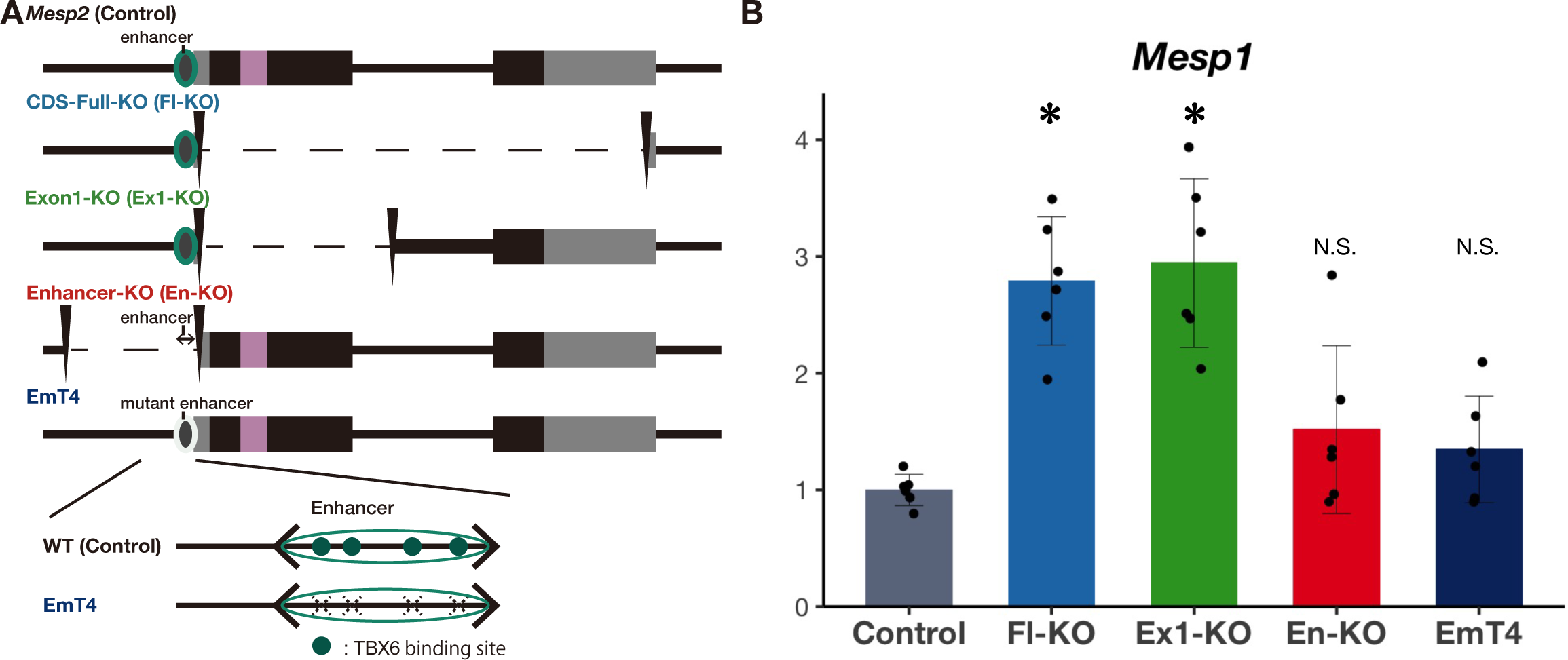
Summary of the conditions of the *Mesp2* locus and compensatory response. A Schematic diagram of *Mesp2* gene structure and its deletions or mutations. B Summary of *Mesp1* expression in *Mesp2* mutant lines. The data of *Mesp1* expression are derived from: control, CDS-Full-KO, and Exon1-KO from Fig 1G; Enhancer-KO from Fig 2D; EmT4 from Fig 2G. These data were normalized by *Mesp1* expression in the control in each experiment. *Asterisk* indicates significant (*p* < 0.05).

**Expanded View Figure 3.**
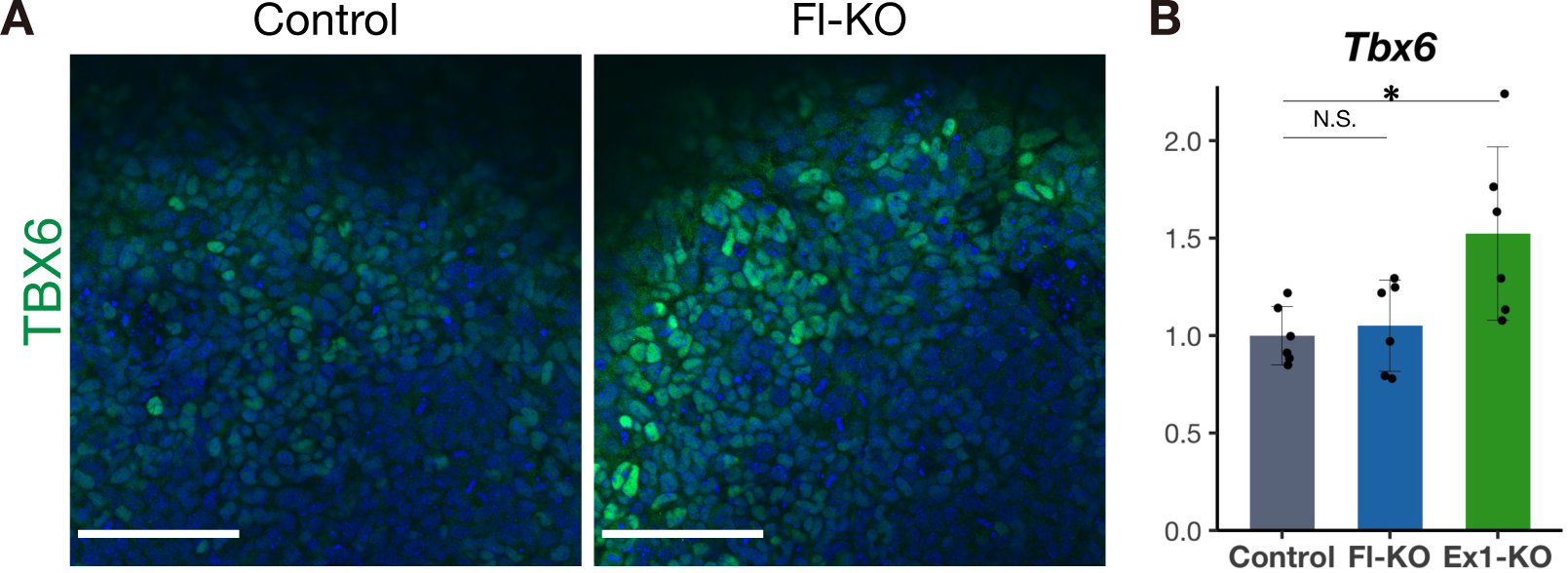
Confirmation of the breakdown of the negative feedback loop by MESP2 loss. A Immunostaining of TBX6 in the *in vitro* PSM of *iTbx6;iNICD*(left) and *iTbx6;iNICD*;*Mesp*2-CDS-Full-KO (right) upon Dox administration. Scale bars: 100 µm. B qPCR analysis of *Tbx6* in *in vitro* PSM (n = 6 cultures for each genotype). *P*-values were calculated by the Mann-Whitney U test comparing *iTbx6;iNICD* and *iTbx6;iNICD*;*Mesp2*-KO lines. Data are presented as the mean ± SD. *Asterisk* indicates significant (p < 0.05).

**Expanded View Figure 4.**
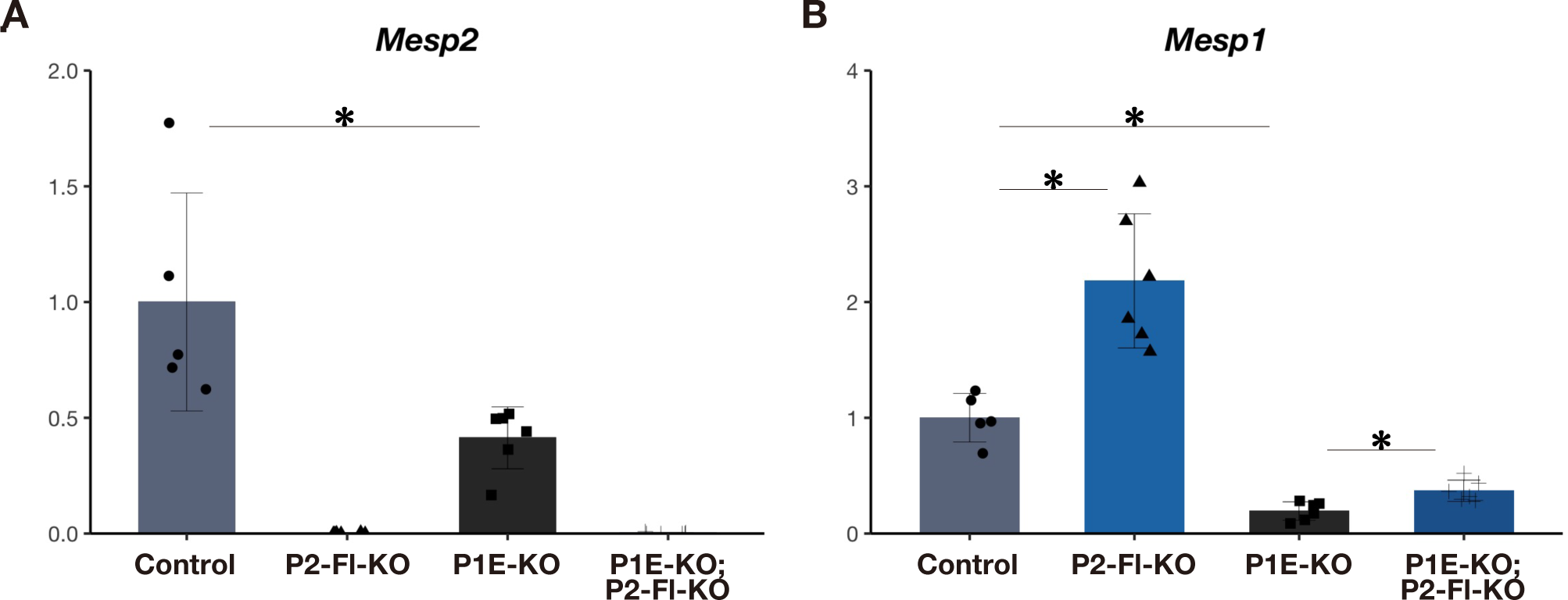
The cooperative function of the *Mesp1*-enhancer in this compensation in another genetic background. A and B qPCR analysis of *Mesp2* (A) and *Mesp1* (B) in *in vitro* PSM; *iTbx6;iNICD*, *iTbx6;iNICD*;*Mesp1*-enhancer-KO, *Mesp2*-CDS-Full-KO with the intact *Mesp1*-enhancer, and additive *Mesp2*-CDS-Full-KO on *iTbx6;iNICD*;*Mesp1*-enhancer-KO (n = 5 or 6 cultures for each genotype). *P*-values were calculated by the Mann-Whitney U test comparing *iTbx6;iNICD*;*Mesp1*-enhancer-KO and others. Data are presented as the mean ± SD. *Asterisk* indicates significant (*p* < 0.05).

**Expanded View Figure 5.**
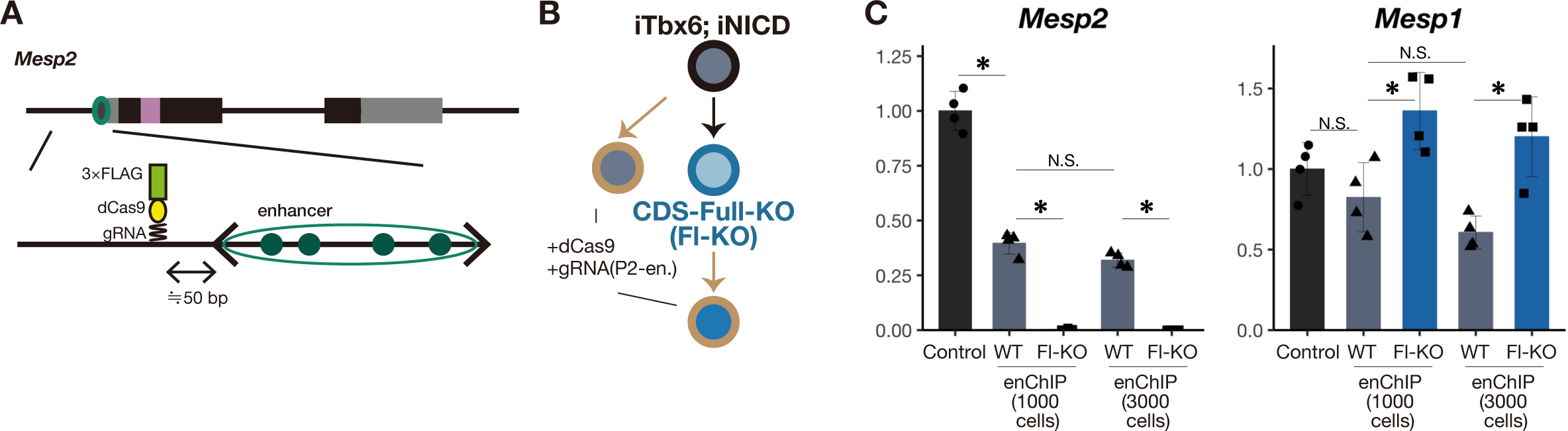
Schematic diagram of enChIP cell lines and the occurrence of compensation between them. A Schematic diagram of the gRNA recognition position proximal to the *Mesp2*-enhancer, which was used for enChIP lines. gRNA sequences are listed in Table 1. B Schematic illustration of the generation of enChIP lines. C qPCR analysis of *Mesp2* and *Mesp1* in the *in vitro* PSM; *iTbx6;iNICD*, and enChIP lines (n = 4 cultures for each genotype). The starting cell number used for PSM induction per well was 1000. For enChIP cell lines, 3000 cells were also used to induce PSM. The expression of *Mesp* genes in PSM from 1000 or 3000 cells was not different. *P*-values were calculated by the Mann-Whitney U test comparing samples indicated in this figure. Data are presented as the mean ± SD. *Asterisk* indicates significant (*p* < 0.05).

**Expanded View Figure 6.**
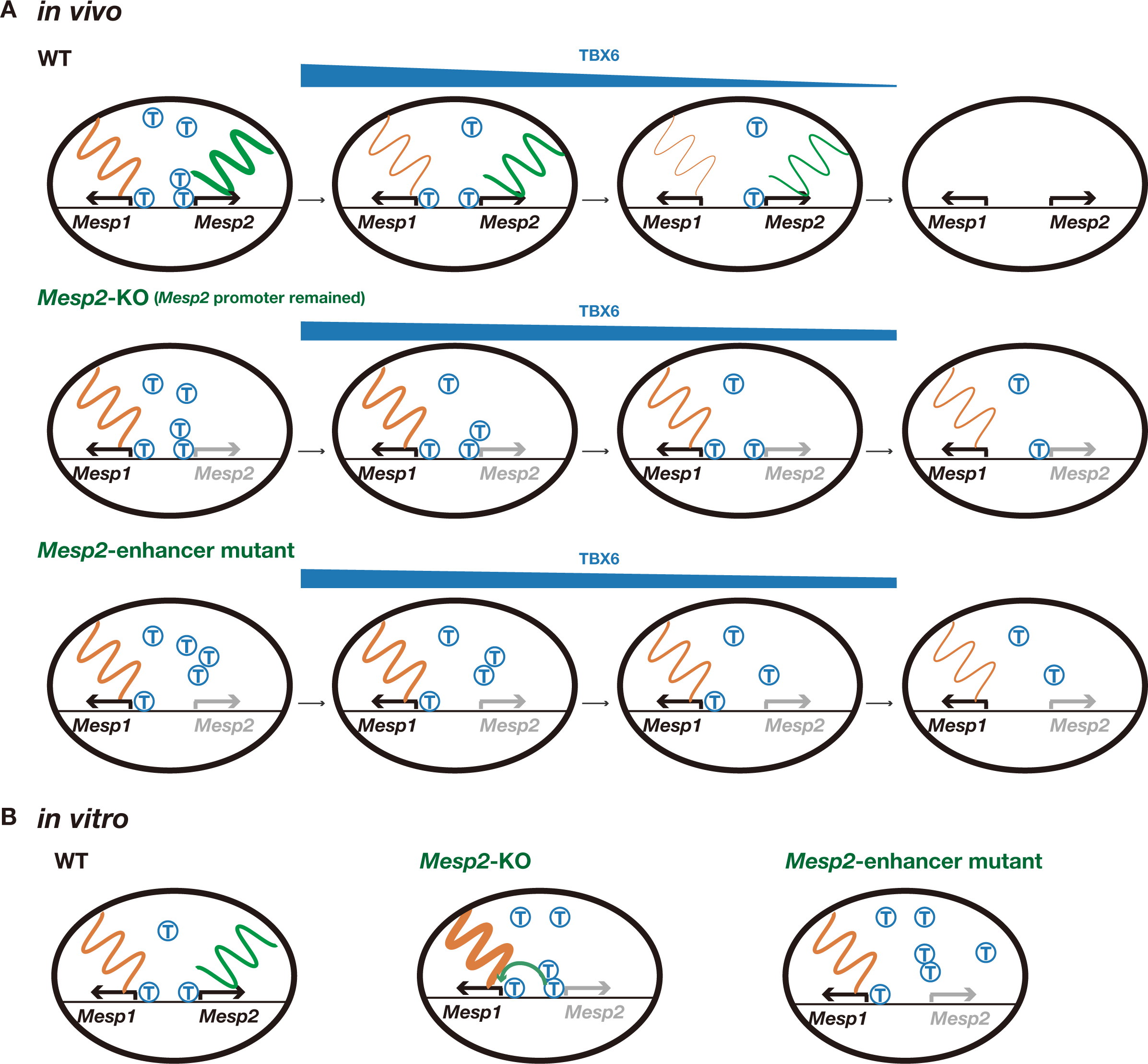
Schematic comparison model of the compensation between *in vitro* and *in vivo Mesp2* mutants. Schematic illustration of the course of *Mesp1* and *Mesp2* expression and TBX6 amount in *in vivo* (A) and *in vitro* (B) PSM. Transcripts of *Mesp1* and *Mesp2* are shown as orange and green wavy lines, respectively, and their thickness indicates the strength of transcription. TBX6 protein is shown as a blue T. Binding sites of T on the line indicate *Mesp1*- or *Mesp2*-enhancer regions. Note that the position of the *Mesp1*-enhancer region is virtually described and not accurate. A Schematic illustration of cells undergoing somitogenesis *in vivo*. TBX6 is gradually degraded by the RIPPLY2-mediated proteasome pathway, and *Mesp1* and *Mesp2* are also gradually degraded in WT. On the other hand, TBX6 remains (Sasaki *et al*, 2011; Zhao *et al*, 2015; Yasuhiko *et al*, 2008) and may continue to bind the *Mesp1*-enhancer in *Mesp2*-KO and *Mesp2*-enhancer mutant mice. The duration of TBX6 binding to the *Mesp1*-enhancer is prolonged, which can increase the amount of *Mesp1* transcripts. B Schematic illustration of cells after the TBX6 and NICD induction *in vitro*. TBX6 is not fully degraded even in the control due to the exogenous induction of TBX6. The TBX6 binding site on the *Mesp1*-enhancer may be occupied in the control and *Mesp2*-KO conditions. The *Mesp1*-enhancer may always be activated and not be different between the control and *Mesp2*-KO conditions.

